# Wound microbiota-mediated correction of matrix metalloproteinase expression promotes re-epithelialization of diabetic wounds

**DOI:** 10.1101/2023.06.30.547263

**Authors:** Ellen K. White, Aayushi Uberoi, Jamie Ting-Chun Pan, Jordan T. Ort, Amy E. Campbell, Sofia M. Murga-Garrido, Jordan C. Harris, Preeti Bhanap, Monica Wei, Nelida Y. Robles, Sue E. Gardner, Elizabeth A. Grice

**Affiliations:** Department of Dermatology, Perelman School of Medicine, University of Pennsylvania, Philadelphia, 19104, USA; College of Nursing, The University of Iowa, Iowa City, 52242, USA

## Abstract

Chronic wounds are a common and costly complication of diabetes, where multifactorial defects contribute to dysregulated skin repair, inflammation, tissue damage, and infection. We previously showed that aspects of the diabetic foot ulcer microbiota were correlated with poor healing outcomes, but many microbial species recovered remain uninvestigated with respect to wound healing. Here we focused on *Alcaligenes faecalis*, a Gram-negative bacterium that is frequently recovered from chronic wounds but rarely causes infection. Treatment of diabetic wounds with *A. faecalis* accelerated healing during early stages. We investigated the underlying mechanisms and found that *A. faecalis* treatment promotes re-epithelialization of diabetic keratinocytes, a process which is necessary for healing but deficient in chronic wounds. Overexpression of matrix metalloproteinases in diabetes contributes to failed epithelialization, and we found that *A. faecalis* treatment balances this overexpression to allow proper healing. This work uncovers a mechanism of bacterial-driven wound repair and provides a foundation for the development of microbiota-based wound interventions.

## Introduction

Skin injury occurs in the context of microbial communities, which can subsequently contaminate and colonize injured tissue. The “wound microbiome” is a ubiquitous component of the wound environment and has been associated with healing outcomes (*1*–*3*). Considered a silent epidemic, chronic wounds affect over six million people in the United States each year (*4*). As the number of people with chronic wounds increases, healthcare costs also surge, with recent estimates of 96 billion spent annually on management of non-healing wounds (*5*). Additionally, patients with chronic wounds experience pain, morbidity, immobility, and even social isolation (*6*–*9*). Given this mounting healthcare threat, there is an unmet need for improved, personalized therapies that target molecular mechanisms specific to wound pathophysiology (*10*). The wound microbiota, and its mechanisms of interacting with host cells during repair, represent a promising source of novel therapeutic targets and/or disease biomarkers.

There is growing evidence that resident skin microbes promote several components of the host skin repair and wound healing responses (*11*). Skin commensal bacteria can promote epithelialization and barrier function by regulating the keratinocyte aryl hydrocarbon receptor (*12,13*). Commensal bacteria can modulate the inflammatory cascade needed for proper repair and regeneration of wounded skin (*14*–*16*). Immune cells that are specific to commensal bacteria are recruited after tissue damage, and can promote injury repair (*17*–*20*). Colonizing skin bacteria have also been shown to promote cutaneous nerve regeneration after injury (*21*). These processes of inflammatory cytokine signaling, immune cell recruitment, epithelialization, and regeneration are dysregulated in chronic wounds. Additionally, dysregulated inflammation, disorganized keratinocyte function, and increased peptidase activity contribute to the wound healing impairment in diabetic skin (*22*). Given the role of skin microbes in promoting diverse tissue repair mechanisms, we hypothesized that microbial-driven responses could be leveraged to correct these dysfunction processes that lead to non-healing wounds.

The risk of infection as well as an incomplete understanding of the wound microbiome has led to clinical practices that seek to eradicate wound microbes. Community-wide culture-independent profiling of the wound microbiota has provided a more comprehensive, less-biased understanding of microbes that are present, as well as their dynamics during healing and complications (*2, 3, 23*). Although well-characterized wound pathogens such as *Staphylococcus aureus* and *Pseudomonas aeruginosa* are commonly detected by these methods, these pathogens do not exist in isolation, but in communities with other microbes that are poorly characterized in the context of cutaneous wounds. We previously conducted shotgun metagenomic sequencing of 195 samples collected from 46 diabetic foot ulcers (DFU), and by rank mean abundance, *S. aureus* and *P. aeruginosa* were the top taxa detected, followed by *Corynebacterium striatum, Propionibacterium* species, and *Alcaligenes faecalis* (*24*). *C. striatum* and *Propionibacterium* spp. are considered human skin commensals, while *A. faecalis* is considered a non-pathogenic environmental bacterium (*25*).

Because commensal microbes can promote many cutaneous repair processes, our objective was to examine the role of wound colonizers and to identify mechanisms of microbial-host crosstalk that contribute to healing outcomes. Here we investigated *Alcaligenes faecalis*, a Gram negative rod that is frequently recovered from chronic wounds by both culture-dependent and independent methods. While shotgun metagenomic sequencing indicated that *A. faecalis* was the 5^th^ most abundant species, it was not associated with clinical outcomes in our DFU cohort. In a murine diabetic wound healing model, we found that *A. faecalis* treatment promoted early wound healing while colonizing the wound bed. *A. faecalis* culture supernatants induced a pro-epithelialization phenotype in diabetic keratinocytes, enhancing migration and proliferation. *A. faecalis* induces this pro-healing phenotype through modulation of matrix metalloproteinase pathways, which are over-active in diabetes and contribute to the highly proteolytic environment that is detrimental to wound healing (*26*–*32*). Thus, we uncover a novel mechanism by which a member of the wound microbiome can promote skin repair and wound healing.

## Results

### Alcaligenes faecalis is a common member of the chronic wound microbiota and promotes early wound closure in a murine diabetic model

We previously identified *A. faecalis* as a prevalent and abundant member of the diabetic foot ulcer (DFU) microbiome, through both culture-dependent and -independent methods (*24*). In a longitudinal prospective cohort study of 100 DFU, we performed quantitative cultures in parallel with culture-independent methods to identify microbial bioburden associated with clinical outcomes (**Fig. 1A**). Cultures identified 44 *A. faecalis* isolates from 14 DFUs (**Fig. 1B**). Shotgun metagenomic sequencing indicated that *A. faecalis* comprised up to 38% mean relative abundance in culture-positive DFUs (**Fig. 1B**). Though *A. faecalis* was ranked 5^th^ by mean relative abundance overall, we did not find an association with DFU outcome in our analysis (*24*). Our findings are supported by additional studies where *Alcaligenes* is consistently identified in chronic wounds other than DFU, including from pressure ulcers (PU), venous leg ulcers (VLU), and sickle cell disease leg ulcers (SCLU) (**Table 1**) (*33*–*40*).

**Table 1:**
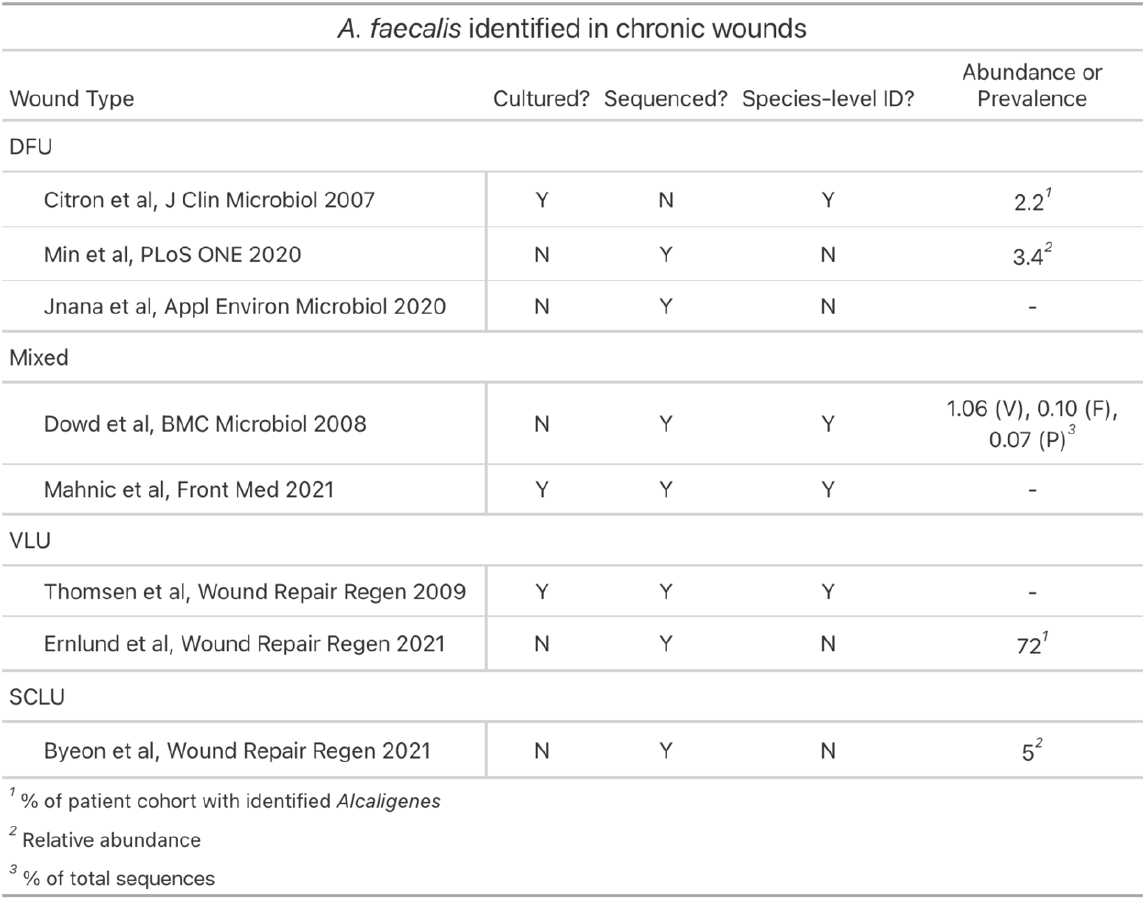
*Alcaligenes* is frequently identified in several chronic wound types. Studies of several chronic wound types including diabetic foot ulcers (DFU), venous leg ulcers (VLU), and sickle cell disease leg ulcers (SCLU) have identified *Alcaligenes* as a prevalent and abundant component of the wound microbiome. Citations by wound type are shown, along with method of *Alcaligenes* identification and whether species-level identification was resolved. Measures of abundance or prevalence are shown in the footnote.

**Fig. 1.**
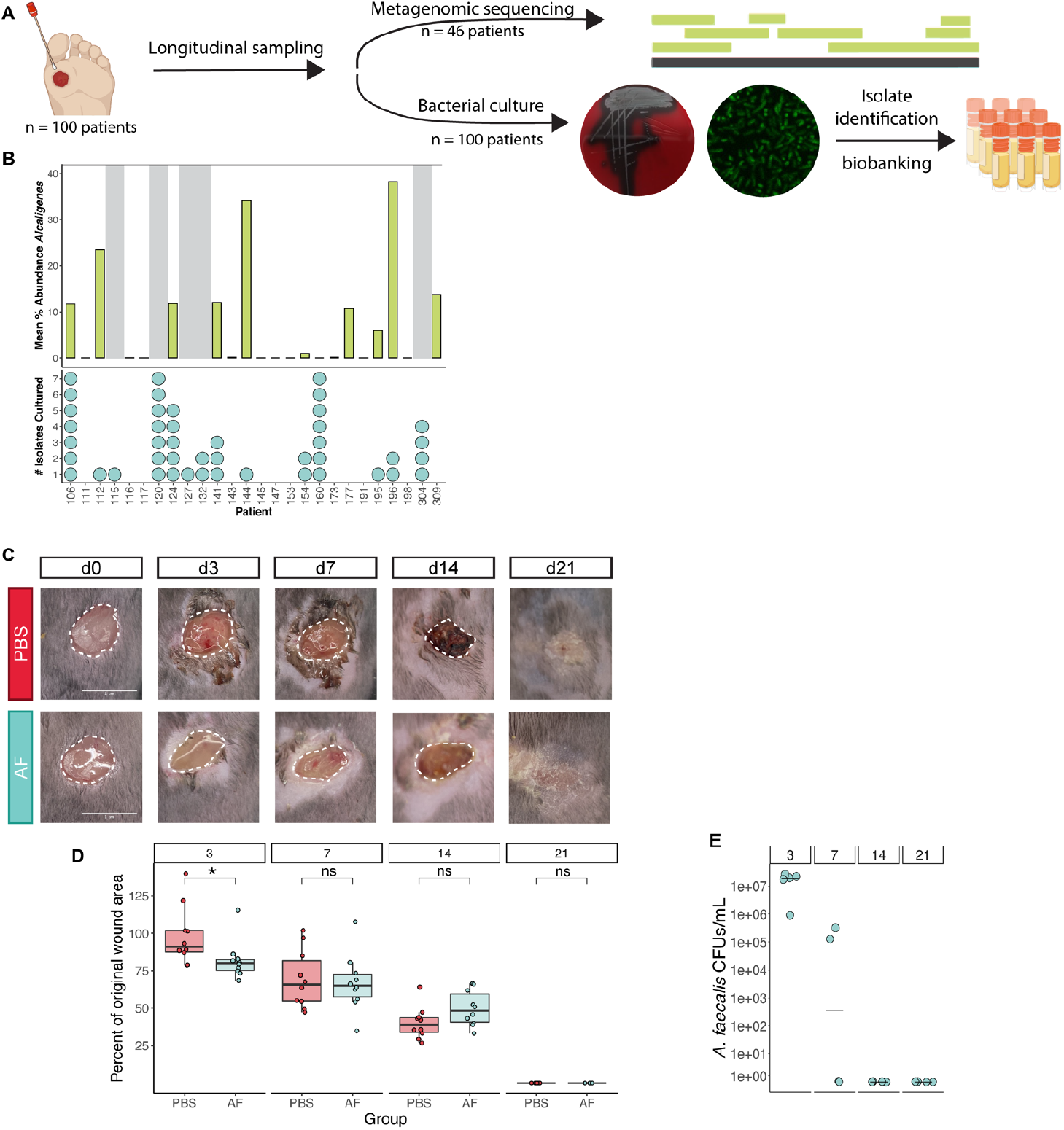
*Alcaligenes faecalis* is prevalent in the wound microbiome and promotes early diabetic wound healing. (**A**) Longitudinal samples collected from diabetic foot ulcers of 100 patients were cultured, and a subset of samples were sent for shotgun metagenomic sequencing. Representative images of *A. faecalis* streaked on blood agar and bacterial fluorescent *in situ* hybridization are shown. (**B**) Prevalence and abundance of *Alcaligenes* across the DFU-100 patient cohort. Top plot shows the mean percent abundance of *Alcaligenes* out of total bacteria for each patient sample’s metagenomic data. Patients that were culture positive but did not have samples sent to metagenomic sequencing performed are greyed out. The number of *A. faecalis* isolates cultured from unique patient samples are displayed in the bottom plot. (**C, D**) 8mm wounds were created in diabetic mice and colonized with vehicle control (PBS) or *A. faecalis* (AF). Longitudinal wound images were taken at days 0, 3, 7, 14, and 21 and wound size (white dashed outline) compared to original wound area was quantified, demonstrating a significant decrease in AF-treated wound size at d3. Scale bar= 1cm; n =5 mice/group, 2 wounds/mouse; * <0.05, p = 0.015 by Wilcoxon rank-sum test; data representative of 2 independent experiments. (**E**) Wound tissue was harvested at each time point for *A. faecalis* colony forming unit (CFU) quantification. n = 4 mice/group (day 3 = 5 mice), 1 wound/mouse.

Despite the lack of association with clinical outcomes, the abundance and prevalence of *A. faecalis* in our cohort motivated us to study the direct impact of *A. faecalis* on diabetic wound healing. To test this, we employed a 12-week old *db/db* diabetic mouse model (B6.BKS(D)-*Lepr*^*db/db*^), which has demonstrated wound healing defects (*41*). We made full-thickness excisional 8mm wounds on the shaved mouse dorsum, treated each wound with 2×10^8^ CFU of a clinical *A. faecalis* isolate or vehicle control, and covered wounds with Tegaderm. Wounds were photographed and wound area was measured at days 0, 3, 7, 14, and 21 (**Fig. 1C**, unedited wound images **Fig. S1A**). *A. faecalis* colonization significantly accelerated wound closure at day 3 compared to vehicle-treated control (**Fig. 1C, D**; p = 0.015 by Wilcoxon rank-sum test). The *A. faecalis*-treated wounds showed no overt signs of infection and maintain well-defined wound margins (**Fig. 1C**). The wound bed appeared healthy despite *A. faecalis* colonization persisting at ∼2×10^7^ CFU at day 3 (**Fig. 1E**). Inflammatory responses during healing promote clearance of bacteria from the wound bed; indeed, partial clearance of *A. faecalis* occurs by day 7 and by day 14 is completely cleared (**Fig. 1E**). Thus, *A. faecalis* colonized the wound bed in parallel to enhanced wound closure during early stages, but does not persist through later stages.

These results of *A. faecalis* treatment are in contrast to treatment of wounds with *Staphylococcus aureus*, a well-established wound pathogen (*42*). Using the same inoculum of a *S. aureus* clinical isolate that delayed healing in diabetic wounds (*43*), wound areas at day 3 were significantly enlarged compared to both vehicle and *A. faecalis*-treated mice (**Fig. S1B;** *: p <= 0.05; **: p <= 0.01; ****: p <= 0.0001 by Wilcoxon rank-sum test). Despite this dichotomous wound healing response, these two bacteria persisted at similar CFU on day 3 post-wounding (**Fig. S1C**; ns = not significant by Wilcoxon rank-sum test). Furthermore, at day 3, *S. aureus* treated wounds have overt signs of inflammation, including erythema, purulent discharge, and macerated wound edges (**Fig. S1C**). Unlike *A. faecalis, S. aureus* is not successfully cleared from the wounds, and these wounds do not heal by the day 21 experimental end point (**Fig. S1D**). These results indicate that unlike pathogenic bacteria, *A. faecalis* can promote early wound closure. Therefore, we sought to understand the mechanisms that underlie this microbially-mediated wound closure.

### A. faecalis improves the epithelialization potential of diabetic keratinocytes

We observed that the strongest effect of *A. faecalis* treatment occurred in the early stages of wound healing, where the critical process of re-epithelialization is initiated. Successful wound closure is definitionally dependent on restoration of the epidermis, or re-epithelialization (*44*). Chronic wounds, including DFUs, are deficient in re-epithelialization as well as the processes of keratinocyte migration and proliferation (*44*–*48*). For these reasons, we investigated the effect of *A. faecalis* on re-epithelialization.

Migration by wound edge keratinocytes is required to close the gap in epithelia that is created by tissue damage and wounding. To examine keratinocyte migration, we modified an *in vitro* scratch wound closure assay to use primary diabetic keratinocytes derived from *db/db* mouse epidermis. Confluent cells were treated with 10% sterile-filtered bacterial conditioned media after creating a “scratch” by removing 2-well tissue culture inserts. Cells were imaged to measure the initial gap and again after 24 hours. *A. faecalis* conditioned media significantly accelerated keratinocyte migration and closed the gap compared to the untreated control (**Fig. 2A**; **: p <= 0.01 by Wilcoxon test). In contrast, cells treated with the same dose of *S. aureus* conditioned media inhibited migration. These differences are not due to changes in keratinocyte viability after 24h incubation with the bacterial conditioned media (**Fig. S2A**).

**Fig. 2.**
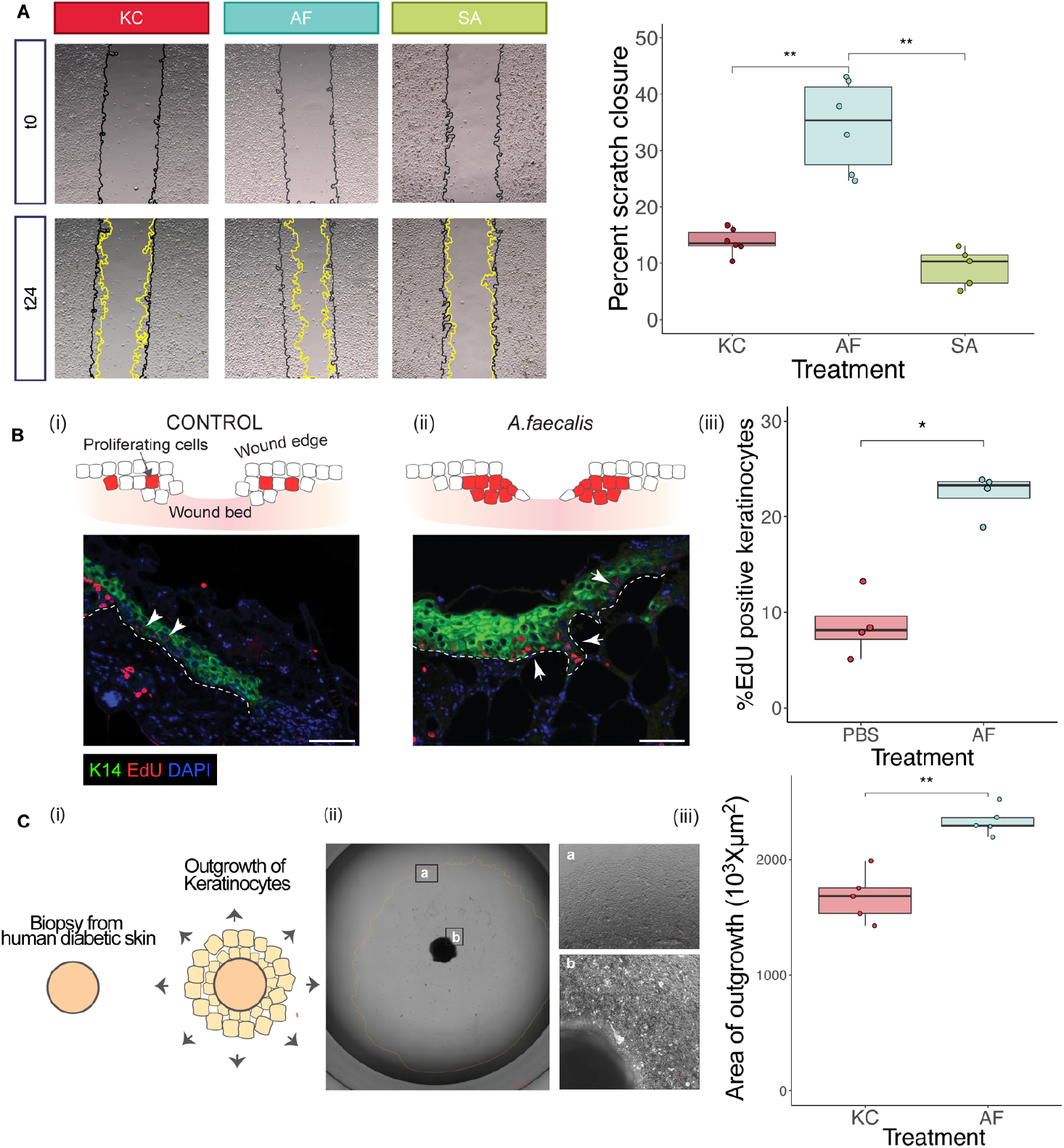
*A. faecalis* enhances diabetic keratinocyte migration and proliferation. (**A**) Gaps were created in diabetic mouse keratinocytes, treated with conditioned media, and *in vitro* wound closure was measured at 24h. Cells were treated with vehicle control (KC), 10% *A. faecalis* conditioned media (AF), or 10% *S. aureus* conditioned media (SA). Initial gap size is drawn in black, and migrating margins are drawn in yellow. **: p <= 0.01 by Wilcoxon test; data representative of >3 independent experiments; n = 6 independent gaps/group. (**B**) 8mm wounds were created in diabetic mice and colonized with PBS (i) or AF (ii). At 3 days post wounding, mice were injected with thymidine analogue 5-ethynyl-2’-deoxyuridine (EdU) 1 hr prior to euthanasia to label proliferating cells. Wound tissue was collected for immunofluorescence staining after paraffin embedding and EdU positive cells (red, white arrows) were detected. Epithelial cells were identified by immunofluorescence staining for Cytokeratin-14 (K14; green) and counterstained with DAPI (blue). Wound edges were imaged (n=4 wounds/condition) and % EdU+ K14+ epithelial cells was determined [number of EdU+ K14+ cells (red nuclei) /total K14+ cells (green cells)] were counted (iii) and compared between PBS and AF treated wounds (*: p <= 0.05 by Wilcoxon test). In Panels (i) and (ii), white dashed lines indicate separation of epithelia from dermis, scale bar (bottom right) =100μm, white arrows indicate examples of EdU+ epithelial cells. (**C**) Ex vivo explant outgrowth assay to measure the epithelialization potential of human diabetic keratinocytes. (i) 1.5 mm biopsies are taken from human diabetic skin, and keratinocyte growth from the explant is quantified. (ii) Representative brightfield image of the explant edge (b) and border of the keratinocyte outgrowth (a) after 10 days of incubation. (iii) Explants were incubated with 10% AF conditioned media or vehicle control (KC), and keratinocyte outgrowth area was quantified at day 10. **: p <= 0.01 by Wilcoxon test; n = 5.

In addition to migration, keratinocytes must also proliferate to replace the damaged epithelia. To examine the effect on keratinocyte proliferation, we wounded *db/db* mice and treated wounds with *A. faecalis* or placebo as described above. At 3 days post-wounding, mice were pulsed with 5-ethynyl-2′-deoxyuridine (EdU), which labels nascent DNA, for 1 hour before euthanasia and wound harvest. Wounds were sectioned and immunostained for cytokeratin 14 (K14) to mark basal keratinocytes and DAPI-stained to mark nuclei. The percent of EdU+ K14+ epithelial cells was determined and compared (**Fig. 2B**). *A. faecalis* treatment significantly increased the percentage of proliferating basal keratinocytes at the wound edge compared to PBS treated control wounds (*: p <= 0.05 by Wilcoxon test). Thus, *A. faecalis* also promotes proliferation of basal keratinocytes in vivo in diabetic wounds.

Keratinocyte outgrowth from skin biopsies is an established *ex vivo* method to assess re-epithelialization potential, and mimics the properties of migration and proliferation exhibited by wound edge keratinocytes (*49*). To test whether *A. faecalis* influences the epithelialization potential of human diabetic skin, we obtained discarded skin from surgery patients with a diabetes diagnosis. Biopsies were created and cultured in 10% *A. faecalis* conditioned media or media alone for 10 days, and keratinocyte outgrowth was quantified. Explants treated with *A. faecalis* conditioned media resulted in significantly larger area of keratinocyte outgrowth (*: p <= 0.05 by Wilcoxon test) compared to vehicle control (**Fig. 2C**). Together these findings demonstrate that *A. faecalis* treatment improved the overall epithelialization potential of keratinocytes, including migration and proliferation, in both human and mouse diabetic skin.

### Matrix-metalloproteinase expression in diabetic wounds is downregulated by *A. faecalis* treatment

Since *A. faecalis* promoted epithelialization processes in diabetic skin, we next investigated the potential mechanisms. To determine how *A. faecalis* treatment influences transcriptional programs in diabetic wounds, we treated murine diabetic wounds with *A. faecalis* as described for **Fig. 1B**, and subsequently harvested wounds after 3 days for RNA extraction and RNAseq (**Fig. 3A**). These wounds were compared to PBS-treated wounds as a negative vehicle control, and *S. aureus*-treated wounds to represent a pathogen-mediated wound healing response.

**Fig. 3.**
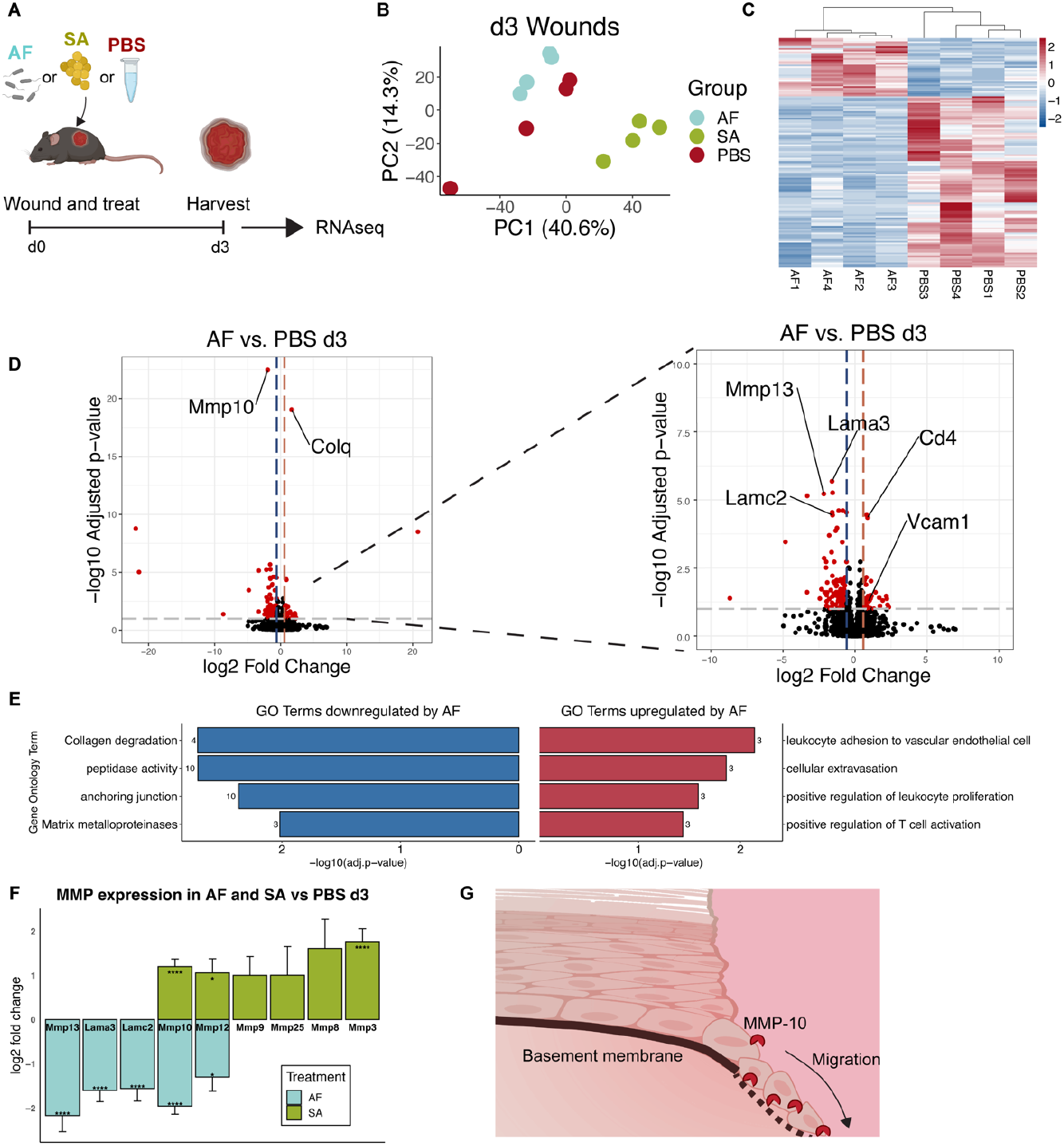
*A. faecalis* treatment decreases MMP expression in murine diabetic wounds. (**A**) Experimental overview for RNA sequencing performed on diabetic murine wounds treated with *A. faecalis* (AF), *S. aureus* (SA), or vehicle control (PBS); n = 4 mice/group; 1 wound/mouse. (**B**) Principal component analysis on day 3 wounds demonstrates distinct clustering of the three treatment groups. (**C**) Heatmap of differential expressed genes (DEG with FDR < 0.01) between the AF and PBS groups at day 3. Unsupervised clustering using Spearman’s correlation was used for hierarchical clustering of the samples, while Pearson’s correlation was used for gene clustering (heatmap rows, dendrogram not shown). Colors are used to codify log fold change values, with red values for increased expression and blue values for decreased expression. (**D**) Volcano plot demonstrating differentially expressed genes in AF vs. PBS wounds at day 3. Horizontal dashed line represents a cut off for the FDR p-adjusted value of <0.1. Vertical lines denote a fold change of ±1.5-fold. Genes that surpass both the p-value and fold change cut off are marked in red, with genes that are downregulated by AF to the left of the plot and upregulated to the right. The inset shows the same volcano plot but at higher resolution. Significant genes of interest are highlighted with gene names on the plot. (**E**) Gene ontology (GO) enrichment was performed on differentially expressed genes for AF vs PBS at day 3. Up and downregulated GO terms of interest are highlighted, with the number of genes matching the term adjacent to the bar plot. (**F**) Gene expression changes of MMPs and related genes in AF vs PBS and SA vs PBS day 3 wounds; * p adj <0.05; **** p adj <0.5 × 10^−4^. (**G**) Conceptual model demonstrating the activity of MMP-10 at the epidermal wound tongue.

Treatment with *A. faecalis* induced a distinct transcriptional profile in day 3 wounds compared to vehicle control or *S. aureus*. (**Fig. 3B**). We performed unsupervised hierarchical clustering of differentially expressed genes of the *A. faecalis* and vehicle-treated groups, demonstrating the modules of up- and down-regulated genes that are distinct in each group (**Fig. 3C**). Treatment with *A. faecalis* led to a greater number of significantly downregulated genes versus upregulated (119 vs. 74, respectively). These results are in contrast to the response of pathogenic *S. aureus* treatment, which resulted in increased expression of many more genes (440) (**Fig. S3A**).

*A. faecalis* significantly upregulated *Colq*, encoding for a subunit of acetylcholinesterase, as well as the immune-related genes *Cd4* and *Vcam1* (**Fig. 3D**). Accordingly, gene ontology (GO) analysis demonstrates that the significantly upregulated gene signatures are related to leukocyte recruitment and T-cell activation (**Fig. 3E**). Of the genes that were downregulated by *A. faecalis*, the most significant were those related to matrix metalloproteinases (MMPs), notably *Mmp10* and its substrates *Lamc2* and *Lama3* (*50, 51*). The downregulated GO pathways were accordingly enriched for signatures related to MMP activity, such as collagen degradation and peptidase activity. In contrast, *S. aureus* treatment led to an increase in GO terms related to peptidase activity, including the gene member *Mmp10*, (**Fig. S3B, C**), along with terms related to cell death, defense responses, and IFN-gamma signatures. Of particular interest is the inverse pattern of MMP-related gene expression. Unlike the significant downregulation of several MMPs induced by *A. faecalis, S. aureus* induces increased expression of six different MMP encoding genes, with MMP-10 being the most significantly different between the two groups (**Fig. 3F**; * p adj <0.05; **** p adj <0.5 × 10^−4^).

During wound healing, MMP-10 is expressed by keratinocytes at the leading edge of the wound and degrades the extracellular-matrix (ECM) components that provide a substrate for keratinocyte migration (*50, 52, 53*) (**Fig. 3G**). However, excessive expression of MMPs, which is promoted by diabetes and hyperglycemia, is deleterious to healing (*28, 54*–*56*). Together, our results suggest that *A. faecalis* promotes diabetic wound healing by inhibiting MMP expression locally to promote re-epithelialization.

### Reduction of excess MMP-10 mediates the pro-epithelialization effects of *A. faecalis*

We focused on MMP-10 as a potential mechanism because it was the most significantly differentially expressed gene and is expressed by keratinocytes at the wound leading edge. To begin to test the hypothesis that *A. faecalis* is moderating MMP-10 levels to promote re-epithelialization, we correlated the RNAseq results with protein expression during diabetic wound healing. We performed immunofluorescence staining for MMP-10 on day 3 wounds (**Fig. 4A**). Consistent with previously reported expression patterns (*52, 53, 57*), MMP-10 was more highly expressed in wound tongue, nearest the wound bed, compared to sparse epidermal staining in sites distal from the wound bed (**Fig. 4B**). MMP-10 protein levels in the proximal wound epidermis trended toward reduction in *A. faecalis*-treated wounds compared to vehicle control (**Fig. 4C**; p = 0.081 by Wilcoxon-rank sum test; sample ImageJ analysis shown in **Fig. S4A**).

**Fig. 4.**
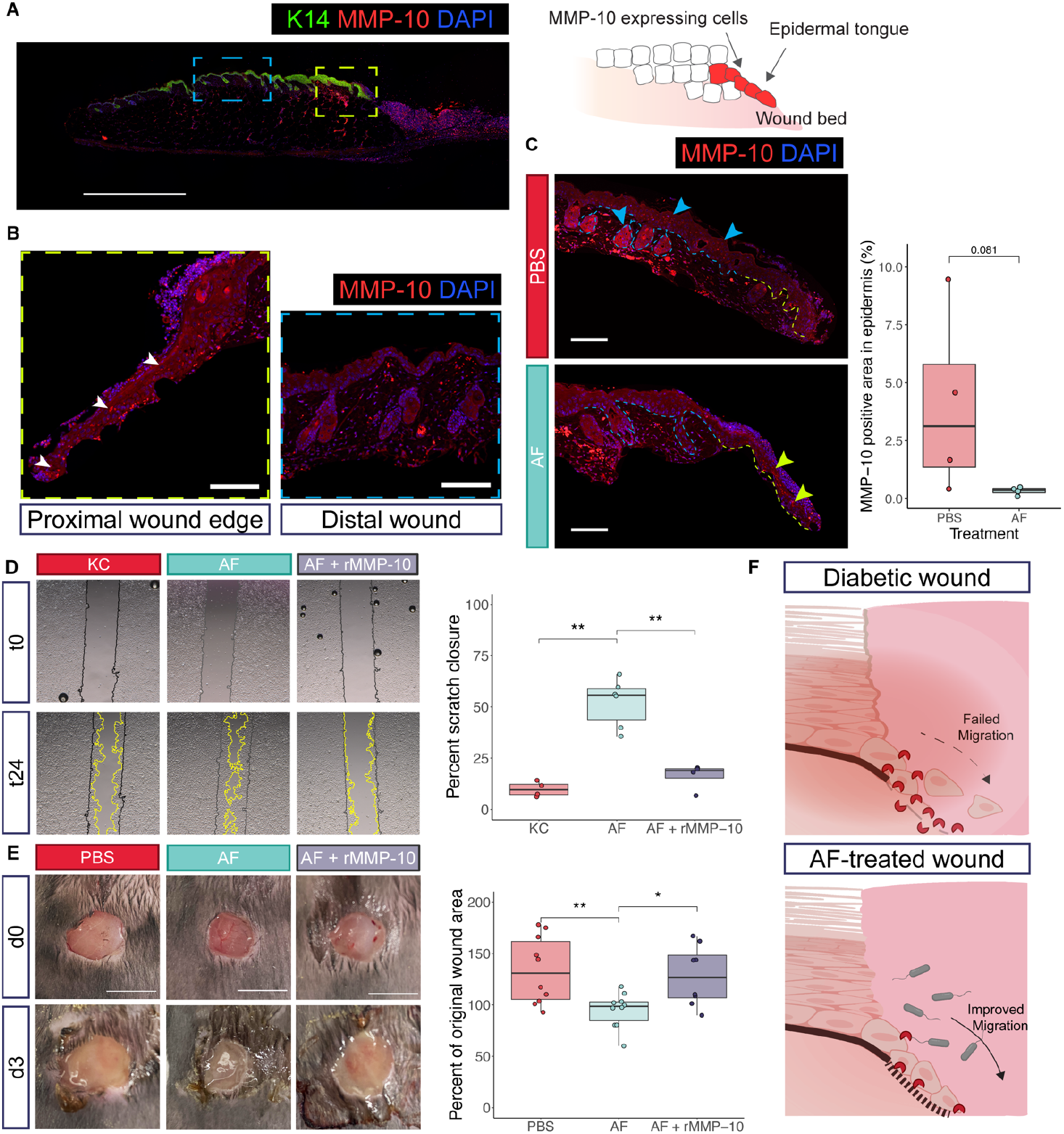
The pro-healing phenotype of *A. faecalis* is dependent on MMP-10 reduction. (**A**) Immunofluorescence staining was performed on paraffin-embedded sagittal sections of day 3 wound tongues. Keratin-14 (K14), MMP-10, and nuclear (DAPI) co-staining were performed. Scale bar = 1000 μm. Blue and green boxes indicate locations of higher magnification insets to show greater detail in (**B**). The positive MMP-10 signal is highlighted by arrowheads in the proximal wound edge, while there is negative MMP-10 staining in the distal wound epidermis. Scale bar = 100 μm. (**C**) The percentage of epidermal area with a positive MMP-10 signal was calculated in the epidermis adjacent to the wound bed. The green dashed line highlights the proximal wound tongue, while the blue dashed line is distal wound epithelium. Arrowheads highlight areas of MMP-10 positive signal. Scale bar = 250 μm, n = 4 mice/group; p = 0.081 by Wilcoxon-rank sum test. (**D**) Gaps were created in diabetic mouse keratinocytes, and *in vitro* wound closure was measured at 24 hours. Initial gap size is drawn in black, and migrating margins are drawn in yellow. AF increases scratch closure, while addition of 200 ng/mL murine recombinant MMP-10 (rMMP-10) abrogates the pro-healing phenotype. [n = 5 independent gaps/group (except n=6 for AF condition); data shown is representative of >3 independent experiments; **: p <= 0.01 by Wilcoxon rank-sum test]. (**E**) 8mm wounds were created in diabetic mice and colonized with PBS, AF, or AF + rMMP-10. Wound images were taken at days 0 and 3 and wound size compared to original wound area was quantified, demonstrating a significant decrease in AF-treated wound size at day 3. Addition of rMMP-10 abolishes AF-mediated accelerated closure. Scale bars = 1 cm; n = 5 mice per group (4 per group for AF + rMMP), and 2 wounds per mouse; *: p <= 0.05; **: p <= 0.01 by Wilcoxon rank-sum test.

Next, we tested the dependence on MMP-10 reduction for the pro-healing effects of *A. faecalis* treatment by performing a scratch assay with murine diabetic keratinocytes as described in **Fig. 2A**. The keratinocytes are seeded on a thin coating of Matrigel, a basement membrane substrate made up of ECM proteins that are digested by MMP-10. To mimic the over-expression of MMP-10 that can occur in diabetic wounds, exogenous recombinant mouse MMP-10 (rMMP-10) was added to *A. faecalis* conditioned media. Addition of excess rMMP-10 significantly abrogates keratinocyte migration observed with *A. faecalis* conditioned media alone (**Fig. 4D;** **: p < 0.01 by Wilcoxon rank-sum test). These differences are not due to changes in keratinocyte viability after 24h incubation with the bacterial conditioned media (**Fig. S2B**).

For *in vivo* validation of these findings, we compared *db/db* wound sizes of *A. faecalis*-treated wounds with those treated with *A. faecalis* plus rMMP-10. As previously observed, *A. faecalis* leads to significantly decreased wound sizes compared to vehicle control at day 3 (**Fig. 4E;** *: p <= 0.05; **: p <= 0.01 by Wilcoxon rank-sum test). However, addition of excess rMMP-10 leads to significantly larger wounds than *A. faecalis* alone, leading to wound sizes that are similar to the vehicle treated control group. Wounds treated with *A. faecalis* and rMMP-10 retained high levels of *A. faecalis* colonization (**Fig. S4B**). Although there was a decrease in *A. faecalis* CFU counts compared to *A. faecalis* alone, this slight decrease is likely due to increased wound size allowing for colonization of other bacterial species, as seen by wound CFU plating. To examine how excess MMP-10 expression affects wound edge morphology, we performed immunohistochemistry on the wound samples, using Masson’s trichrome stain to distinguish the epidermal tongue from the underlying dermis (**Fig. S4C**). Consistent with wound size quantification, *A. faecalis*-treated wounds were initiating re-epithelialization, as evidenced by visible wound tongues. However, the vehicle and rMMP-10-treated groups had little keratinocyte migration at the wound edge. Together, these results indicate that *A. faecalis* promotes early diabetic wound closure by decreasing rMMP-10 overexpression (**Fig. 4F**).

## Discussion

Improved therapeutic approaches are needed for non-healing wounds, as they present a major challenge to the healthcare system by increasing treatment costs as well as rates of morbidity and mortality. The skin microbiome exists at the interface of all cutaneous wounds, but its potential as a novel therapeutic target remains untapped. We previously deeply characterized the microbiome of DFU by shotgun metagenomic sequencing and culture-based approaches. In addition to wound pathogens and skin-resident commensals, we also detected microbes that have not been studied in the context of wound healing. Here, we focused on determining the functional relevance of what was previously considered a wound “bystander” or contaminant, *A. faecalis*. Our findings show that *A. faecalis* treatment promotes diabetic wound closure by enhancing the epithelialization potential of diabetic keratinocytes. *A. faecalis* treatment of diabetic wounds downregulated expression of MMPs, which are overexpressed in diabetic skin and contribute to delayed wound repair. Thus our results suggest that leveraging selective interactions between microbiota and host could target specific process that are dysregulated during diabetic wound healing.

*A. faecalis* has been previously characterized as an environmental bacteria found in water and soil, and rarely causes infection (*25, 58*). The consistent detection of *A. faecalis* through both culture and sequencing-based methods, with some patients being colonized across multiple time points, indicates that *A. faecalis* is likely a wound inhabitant rather than transient contaminant. In line with this hypothesis, *A. faecalis* has been detected in multiple wound types across the globe (*33*–*40*). It has been referred to as a member of the “core microbiome” in DFUs (*35*) and noted as an abundant, understudied bacteria “that should be considered in future studies” (*34*). Here we show that *A. faecalis* could play a more significant role in wound healing by enhancing re-epithelialization and inhibiting aberrant expression of proteases that are detrimental to wound healing. These results demonstrate the importance of looking beyond wound pathogens in the pursuit of actionable therapeutic targets in the wound microbiome.

It is well established that MMPs are dysregulated in diabetic healing (*26*), whereby MMPs are overexpressed and over activated, along with a concomitant decrease in their inhibitors (*27*– *29*). This imbalance has been shown to inhibit proper keratinocyte migration and delay healing (*30*–*32*). We focused specifically on MMP-10, or stromelysin-2, as it was the most significantly decreased gene in the setting of *A. faecalis* treatment and was upregulated by the pathogen *S. aureus*. Under normal healing conditions, MMP-10 facilitates re-epithelialization by cleaving hemidesmosome and desmosome components to release keratinocytes both from the ECM as well as from cell-to-cell adhesions, respectively (*51*). It is typically only expressed in wounded keratinocytes and exclusively localized to the migrating tip of the epidermal tongue (*52, 53, 57*). Furthermore, this peptidase has been shown to be increased diabetic skin (*53*) and other diabetic epithelium (*59*). Increased expression of MMP-10 has been shown to inhibit re-epithelialization (*57*). Constitutive overexpression of MMP-10 leads to an abnormal migrating epidermal tip with “scattered” keratinocytes, inappropriate keratinocyte contact with ECM components, as well as apoptotic keratinocytes in the wound tongue (*50*). Although our studies focus on the effects of MMP-10 in the wounded epidermis, it is possible that MMP-10 overexpression may be affecting other wound healing compartments. For example, MMP-10 has been shown to regulate resident macrophage collagenase activity in the skin (*60*). Given the importance of temporal, spatial, and cellular dynamics for MMP signaling, future investigations will take advantage of high-resolution methods to parse apart these dynamics.

MMP activity, like any inflammatory process, needs to be finely-tuned and balanced during healing (*61*), including a well-orchestrated temporal and spatial localization of MMP expression (*62, 63*). This organization is disrupted in diabetic wounds, which are considered “stuck” in the early inflammatory phase, and unable to progress to the proliferative phase (*26*). Therefore, by rebalancing MMP-10 expression, *A. faecalis* may be able to push diabetic wounds towards a pro-epithelialization environment. This work may also have implications beyond DFUs, as MMPs are upregulated in pressure ulcers and venous leg ulcers (*64, 65*), which are also commonly colonized by *A. faecalis* (**Table 1**).

Future work will be needed to delineate the pathway by which *A. faecalis* modulates keratinocyte MMP expression to promote epithelialization. Keratinocytes are the first responders of the innate immune system to skin barrier breach. As such, they are equipped with myriad receptors that detect bacterial products. There is growing appreciation for injury signals, beyond pathogen- and damage-associated molecular patterns (*66*), that are detected by keratinocytes and can induce re-epithelialization (*67*). It is reasonable to hypothesize that an *A. faecalis*-derived signal is sensed by keratinocyte receptors and induces a pro-migratory phenotype, though the identity of that signal is unknown at this time.

We did not find any correlative evidence from human studies to suggest that *A. faecalis* influences clinical outcomes of wound healing. This could be due to insufficiently powered studies, or alternatively, biological factors that limit the effect of a single microbe in a greater network of microbe-microbe and host-microbe interactions. Our reductionist approach was valuable in delineating the functional potential of *A. faecalis*. Expanding these findings to a community-wide framework will be crucial to understanding how pro-healing “commensals” interact with pathogens and other members of the wound microbiome. These interactions will need to be considered for the success of any microbial-directed therapy for wound healing.

Our goal was to investigate the “dark matter” of the wound microbiome and resulted in identification of *A. faecalis* as a wound resident with the potential to correct aberrant processes that lead to delayed healing. These results reveal a novel therapeutic target and underscore why the wound microbiome, beyond well-established pathogens, should be considered. In light of broad-spectrum antibiotic therapies that deplete commensal microbes such as *A. faecalis* while selecting for pathogen resistance and virulence, our work underscores the need for judicial antibiotic prescription for chronic wounds, especially in the absence of clinical signs of infection (*68*). Further dissection of these complex host-microbiome dynamics during tissue repair, of *A. faecalis* and other members of the wound microbiota, should reveal additional promising and novel therapeutic targets for chronic wounds.

## Materials and Methods

### Culture- and sequencing-based detection of *A. faecalis* in DFUs

Longitudinal samples for microbial analysis were collected from DFUs of 100 patients as previously described (*69, 70*). Swabs were collected using the Levine technique to sample deep tissue fluid (*71*), resuspended in tryptic soy broth (TSB), and plated on blood agar, eosin-methylene blue agar, and Chromagar plates. *A. faecalis* isolates were identified from eosin-methylene blue agar plates by morphology and gram staining for gram-negative rods, using standard microbiological procedures (*72*). The isolates were further identified by Sanger sequencing of the 16S ribosomal RNA gene and matrix-assisted laser desorption ionization-time of flight (MALDI-TOF) mass spectrometry at the Pennsylvania Animal Diagnostic Laboratory System. Metagenomic sequencing was performed on a subset of the DFU specimens as previously described (*24*). Alcaligenes abundance was estimated using MetaPhlAn2 with default parameters and is reported as the mean % of bacterial abundance across samples from each patient (*73*).

### Bacterial strains and preparation

The *A. faecalis* (EGM#4-61) and *S. aureus* (EGM#5-76) strains were cultured as described above and maintained in the Grice lab culture repository (number in parenthesis indicates unique lab identifier code). Bacteria were prepared for wounding treatment by streaking onto blood agar and incubating at 37°C for 24 hours. Single colonies were then used to inoculate TSB and then grown by shaking at 250rpm at 37°C for 18 hours. The bacteria culture was then spun down at 13,000 rpm for 2 minutes. To prepare a glycerol stock, the pellet was resuspended in PBS with 10% glycerol then kept at -80°C until the day of wounding. At the same time, a portion of the overnight culture was serially diluted and plated for CFU calculation.

For conditioned media preparation, the strains were streaked out on blood agar and grown in TSB as described above. The TSB bacterial culture was then used to inoculate human keratinocyte media without antibiotics and grown by shaking at 250 rpm at 37°C for 24 hours. The bacteria culture was then spun down at 4,000 rpm for 30 minutes at 4°C. The supernatant was passed through a 0.22 μm filter for sterilization. At the same time, a portion of the overnight culture was serially diluted and plated for CFU calculation.

### Animal models and husbandry conditions

All mouse experiments were conducted under protocols approved by the University of Pennsylvania Institutional Animal Care and Use Committee (Protocol 804065). All mice were housed and maintained in an ABSL II and specific pathogen free facility in the Clinical Research Building vivarium at the University of Pennsylvania. The following strain of mice was used in these studies: B6.BKS(D)-*Lepr*^*db/db*^/J (JAX stock #000697) (*41*).

### Excisional wounding and bacterial colonization of murine diabetic wounds

One day prior to wounding, the mouse dorsum was shaved with an electric razor. Mice were singly housed on the day of wounding. Pain control was administered with extended-release buprenorphine and bupivacaine injections, and isoflurane was administered for anesthesia. Two full thickness excisional wounds were made on each mouse back using an 8mm punch biopsy tool (Miltek). For bacterial treatment of wounds, 2×10^8^ CFU of *A. faecalis* or *S. aureus* in 10 μL of PBS was applied to each wound. For vehicle-control treatment, the same volume of PBS with 10% glycerol was applied to the wounds. All wounds were photographed and measured at time of wounding (day 0), and subsequent time points (e.g. day 3). After day 0 imaging, the wounds were covered with Tegaderm (3M) to prevent external bacterial contamination. Tegaderm was left on until day 3 wound imaging, and if the experimental timeline continued, wounds were re-covered with Tegaderm until day 7. Wounds were blinded, then quantified in ImageJ (version 1.53k) using a ruler-calibrated measurement, with each wound being measured 3 independent times.

For addition of exogenous MMP-10 to the wounds, murine recombinant MMP-10 (RayBiotech 230-00748-10) was diluted in PBS and 4 ug was added to each wound. This dose was scaled to a similar molarity as previous rMMP treatments of wounds (*74*). For protein and vehicle control, purified BSA (Sigma) was diluted in PBS to the same molarity as rMMP-10, and then added to the PBS control and *A. faecalis* treatment.

### Bacterial CFU quantification of wounds

For bacterial quantification from wound tissue, a 12 mm punch biopsy tool (Miltek) was used to dissect the wound bed and surrounding wound edge tissue from the back skin. Wound tissue was homogenized with horizontal vortexing using ceramic beads (MP). The homogenate was serially diluted, and each dilution was plated in duplicate on blood agar plates to quantify CFUs. For bacterial quantification by swabbing, sterile cotton tipped applicators (Puritan) were dipped in PBS and used to swab the wound bed using the Levine technique (*71*). Swabs were collected in PBS, and the PBS was subsequently serially diluted and plated on blood agar as described above.

### Murine diabetic keratinocyte derivation

Primary keratinocytes were derived from tail skin of adult *db/db* diabetic mice [B6.BKS(D)-*Lepr*^*db/db*^] mice as previously described (*75*). Briefly, skin was dissected from the tail bone, and incubated in 4U/mL dispase (Sigma) overnight at 4°C. The epidermis was then separated from the dermis, and the epidermis was incubated with 0.05% Trypsin-EDTA at room temperature for 20 minutes. The digestion reaction was then quenched with DMEM + 5% FBS. A single-cell suspension was generated by vigorously rubbing the epidermal sheet against a Petri dish. The cell suspension was then passed through a 100 μm filter, centrifuged at 4°C for 5 minutes at 180*g*. For plating, the cell pellet was resuspended in cold low Ca^2+^ keratinocyte serum free media (Life Tech) supplemented with bovine pituitary extract, human epidermal growth factor, 1% FBS, 4% DMEM, and antibiotics-antimycotics.

### Scratch closure assay

Prior to keratinocyte seeding, cell culture plates were thin-coated with Matrigel (Corning), an ECM equivalent. Two-well inserts that create a defined cell-free gap (Ibidi) were placed in a 24-well plate. Primary diabetic keratinocytes (derivation described above) were seeded in the inserts at a density to reach 100% confluence the following day. After overnight incubation, the inserts were removed, cells were washed and then treated with 10% bacterial conditioned media or vehicle control (10% keratinocyte media used for bacterial growth). The starting wound gap was immediately imaged on the Keyence BZ-X710 (t=0 hours). After 24 hour incubation at 37°C and 5% CO_2_, the identical gap location was imaged (t=24 hours). Microscope images were blinded, and gap area at t=0 and t=24 was quantified using the ImageJ “Wound healing size tool, updated” (*76*). For addition of exogenous MMP-10 to the scratches, murine recombinant MMP-10 (RayBiotech 230-00748-10) was diluted in keratinocyte media to the indicated concentrations.

### Viability of KCs

The alamarBlue reagent (Invitrogen) was used to determine viability of primary diabetic keratinocytes. At the experimental endpoint, alamarBlue was diluted 1:10 in keratinocyte media, incubated for 4-6 hours, and the supernatant fluorescence was read with excitation at 560 nm and emission at 600 nm. Fluorescence of alamarBlue added to keratinocyte media alone was subtracted as background.

### EdU incorporation, detection, and visualization

A protocol to detect proliferating cells in wounds was used to measure incorporation of modified thymidine analogue 5-ethynyl-2’-deoxyuridine (*77*). Mice were injected with EdU (Abcam, #ab146186) at 25μg/gm weight of mice 1 hour prior to euthanasia. Tissue was collected and fixed in 4% Paraformaldehyde, paraffin embedded and sectioned as described in subsequent sections. Tissue was deparaffinized (2 times in Xylene Substitute, Sigma Aldrich, Cat #A5597), hydrated after series of ethanol graded rinses (1 time each in 100% Ethanol, 95% Ethanol, 70% Ethanol, 50% Ethanol, respectively), and finally rinsed in 1X PBS, 3 times. Tissue was permeabilized in 1X PBS containing 0.5% Triton100 for 1 hour and incubated in EdU detection cocktail. The EdU cocktail is made freshly each time and consists of: 100mM TBS (pH=7.6), 4mM Copper Sulphate, 100mM Sodium Ascorbate and 5μM Sulfo-Cyanine Azide (Lumiprobe Corporation, Cat #B1330). Tissue was rinsed 3 times in 1X PBS and then incubated with 1:5000 dilution rabbit polyclonal anti-Keratin-14 antibody (Biolegend, Clone Poly19053) for 1 hour and then detected with 1:10000 dilution goat anti-Rabbit IgG Secondary Antibody conjugated with Alexa Fluor™ 488 (ThermoFisher Scientific, Cat #A-11008). Tissue was counterstained with DAPI, cover-slipped and imaged. Wide-field fluorescent images were acquired using a 20X lens objective on a Leica DM6000 Widefield Fluorescence Microscope at the University of Pennsylvania, School of Veterinary Medicine Imaging core. Total EdU positive epithelial cells (Krt14 positive cells that were also EdU positive) were counted manually on the composite merged images using ImageJ cell counter feature.

### Outgrowth assay

The explant keratinocyte outgrowth assay was modified as described previously (*49, 78*). Diabetic skin discarded from Mohs surgery was obtained, and subcutaneous fat was removed with a scalpel. Explants were excised using 1.5mm dermal biopsy punch (Miltex) and adhered to the bottom of a 48-well tissue culture dish coated with collagen. The explants were secured using 2 μl Matrigel and submerged in growth medium consisting of Keratinocyte Serum Free Media (KSFM) with keratinocyte growth supplements (Life Technologies #17005042) and Medium 154 (Life Technologies #M154500). Explants were treated with 10% *A. faecalis* conditioned media. Outgrowths from explants were imaged at 10 days post establishment on a Keyence BZ-X710 microscope (Plan Fluor 10X lens) equipped with a motorized stage. The images were stitched using the inbuilt algorithm accompanying the BZ-X-wide image viewer and area of outgrowth was measured using ImageJ.

### MMP-10 immunofluorescence sample preparation, detection, and visualization

Murine wound tissue was collected, fixed overnight in 4% paraformaldehyde, then paraffin embedded and sectioned by the Cutaneous Phenomics and Transcriptomics (CPAT) core of the Penn Skin Biology and Diseases Resource-based Center (SBDRC). Immunofluorescence staining was also performed by the CPAT core. Tissue sections were deparaffinized and rehydrated as described agove. Heat-induced antigen retrieval was performed in 0.01M citrate buffer, pH 6.0 and blocked in 10% normal goat serum. The following antibodies were used for IF staining: K14 1:500 (Biolegend 906004) with secondary Alexaflour 488 conjugated goat anti-chicken 1:500 (Invitrogen A-11039); and MMP-10, 2.0 μg/mL (Invitrogen PA5-79677), with secondary Alexaflour 555 conjugated goat anti-rabbit 1:1000 (Invitrogen A-21428). Primary antibodies were incubated overnight at 4°C.

Fluorescent images were taken on the Leica DM6B-Z fluorescent microscope using the 20X lens objective. For the MMP-10 wound imaging, the light exposure for K14 and DAPI channels was 150 ms, and 125 ms for the MMP-10 channel. Tiled images were stitched together in the LAS-X software (version 3.7.6.25997) then prepared for export using Fiji (version 2.9.0) and processed in Adobe Photoshop (version 24.1.1). MMP-10 signal quantification was performed in ImageJ (version 1.53k); all images were blinded and global thresholding was performed to the same value cutoff on all images to distinguish positive vs. negative MMP-10 signal. A length along the epidermis equal to 1000 μm from the tip of the epidermal wound tongue was measured, and the epidermis of this segment was selected using the freehand tool. The total area was measured, and then the area of positive signal above the threshold was measured to calculate positive MMP-10 signal in the epidermis.

### RNA-sequencing of wound tissue

#### Sample preparation and sequencing

To harvest wounds for RNAseq, a 12 mm punch biopsy tool (Miltek) was used to dissect the wound bed and surrounding wound edge tissue. Tissue was then immediately snap frozen on liquid nitrogen. For RNA extraction, tissue was first cryo-pulverized using the CP02 CryoPrep automated dry pulverizer (Covaris). Trizol (Invitrogen) based phenol-chloroform extraction was then performed for RNA purification, followed by on-column DNase digestion (Qiagen). RNA integrity was assessed with a 2100 Bioanalyzer and the Eukaryote Total RNA 6000 Pico Kit (Agilent). All but one sample had an RNA Integrity Number (RIN) >6. RNA concentration was quantified on Qubit using the RNA Broad Range assay (Invitrogen). Library preparation and sequencing was performed by the Children’s Hospital of Philadelphia High Throughput Sequencing Core. Ribosomal RNA was depleted and libraries were constructed using the Illumina Stranded Total RNA Prep with Ribo-Zero Plus (Illumina) using 100 ng of RNA input. IDT for Illumina UD indexes were ligated to the libraries. Library quality control was performed, and then libraries were sequenced on the NovaSeq 6000 using the S2 flow cell, generating 50-bp paired-end reads.

#### Analysis

Quality control was performed on the sequencing reads using FastQC (version 0.12.1) (*79*). Reads were mapped to the reference genome [Genome Reference Consortium Mouse Build 39 (GRCm39)] using kallisto (version 0.48.0) (*80*). Subsequent analysis was performed in the R computing environment (version 4.1.3) in RStudio (version 2022.02.1+461). Transcript-level abundances were imported with tximport (version 1.22.0) using the length scaled transcripts per million (*81*). Differential gene expression analysis was performed using the DESeq2 package (version 1.34.0) (*82*). Variance-stabilized data was used for principal component analysis. Pairwise differentially expressed genes (DEGs) between treatments were obtained by setting corresponding ‘‘contrasts’’ for each pairwise comparison. The Benjamini-Hochberg (B-H) procedure was used as a multiple-testing correction for adjusted p-values (*83*). Gene ontology (GO) analysis was performed using the gprofiler2 package (version 0.2.1) and the B-H FDR correction method was used to adjust the enrichment p-values (*84*).

### Trichrome staining and visualization

Wound sections were prepared, embedded, and de-paraffinized as described above. Masson’s Trichrome staining was performed by the CPAT core of the SBDRC. Brightfield images were taken on the Keyence BZ-X710 microscope (Plan Fluor 10X lens).

### Statistical analysis

Statistical analyses were performed with the R computing environment (version 4.1.3) and ggplot2 (version 3.3.6) in RStudio (version 2022.02.1+461). For boxplots, the center line shows the median value, while the box limits represent the interquartile range (IQR), and whiskers extend up to 1.5 times the IQR. To compare wound sizes of two independent treatment groups from the excisional wounding experiments at a given time point, we performed the Wilcoxon rank-sum test. Similarly, gap sizes of two independent treatment groups in the *in vitro* scratch closure assay were compared with the Wilcoxon rank-sum test. The Wilcoxon rank-sum test was also used to compare immunofluorescence quantification between two independent treatment groups. For RNAseq analysis, p values were adjusted using the Bonferroni-Hochberg method. P values and sample sizes are shown in main text and figure legends.

## Acknowledgments

We would like to thank the Penn Skin Biology and Diseases Resource-based Center CPAT and STAR cores for their invaluable assistance with histopathology and provision of skin samples. We would also like to thank the Children’s Hospital of Philadelphia High Throughput Sequencing Core for their support of the RNAseq experiment. We are very grateful to current and former members of the Grice lab and the Department of Dermatology for critical discussion and review of the work. We would like to thank the clinical staff and the University of Pennsylvania and University of Iowa for assistance with patient sample collection, as well as the patients who generously participated in the study. We thank Denis Lee, McArdle Laboratory for Cancer Research, University of Wisconsin-Madison for sharing details of home-based protocol for measuring EdU incorporation. Parts of the conceptual figures were created with BioRender.com.

## Funding

NIH National Institute of Nursing Research R01NR009448 (SEG)

NIH National Institute of Nursing Research R01NR015639 (EAG)

NIH National Institute of Arthritis and Musculoskeletal and Skin Diseases P30AR069589 (EAG)

NIH National Institute of Arthritis and Musculoskeletal and Skin Diseases F31AR079852 (EKW)

NIH National Institute of Arthritis and Musculoskeletal and Skin Diseases T32AR007465 (EKW, JCH)

Prevent Cancer Foundation Awesome Games Done Quick fellowship (AU)

NIH National Institute of Arthritis and Musculoskeletal and Skin Diseases K99 AR081404 (AU)

Penn SBDRC Pilot & Feasibility Grant P30AR069589 (AU)

Penn Blavatnik Family Fellowship (AC)

NIH National Institute of Arthritis and Musculoskeletal and Skin Diseases F31AR079845 (JCH)

NIH National Institute of Allergy and Infectious Diseases 5T32AI141393 (MW)

NIH National Institute of General Medical Sciences R25 GM071745-19 (NYR)

## Author contributions

Conceptualization: EKW, EAG

Methodology: EKW, AU, JP, JTO, AEC, JCH

Investigation: EKW, AU, JP, JTO, AEC, SMG, JCH, PB, MW, NYR

Visualization: EKW, AU, AEC

Supervision: EAG Writing—original draft: EKW

Writing—review & editing: EKW, AU, SMG, AEC, EAG, SEG

## Data and materials availability

All data are available in the main text or the supplementary materials. Upon acceptance, the RNA sequencing dataset generated and analyzed in this study will be publicly available for download in the NCBI Gene Expression Omnibus (GEO). Code is available upon request.

## Supplementary Materials

### This PDF file includes

Figs. S1 to S4

**Fig. S1.**
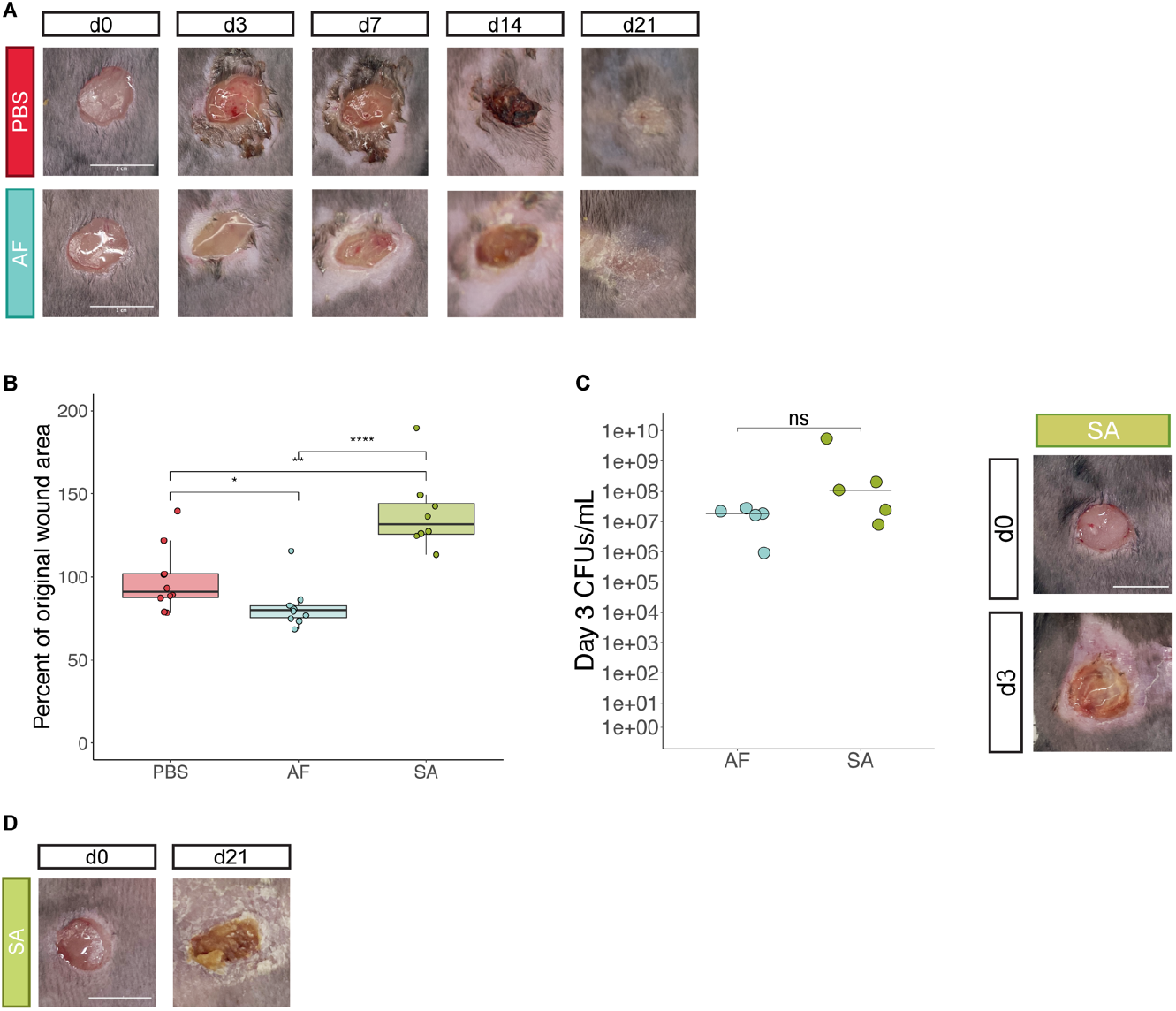
*A. faecalis* has high levels of colonization in day 3 wounds, but does not induce a pathogenic response. (**A**) Wounds were created and imaged as in **Fig. 1C**.; raw images that do not have wound area outlines are shown. (**B**) As in **Fig. 1C**,**D**, diabetic wounds were colonized with a DFU-100 isolate of *S. aureus* (SA). Day 3 wound quantification demonstrates significantly larger wound size of SA-treated wounds; n =4 mice/group, 2 wounds/mouse; *: p <= 0.05; **: p <= 0.01; ****: p <= 0.0001 by Wilcoxon rank-sum test; data representative of 2 independent experiments. (**C**) SA-treated wounds have larger wound area and worse gross pathology compared to AF-treated wounds despite similar levels of d3 SA CFUs vs AF CFUs; n=5 mice/group, 1 wound/mouse; ns = not significant by Wilcoxon rank-sum test. (**D**) Representative image of day 21 wound treated with SA as described in **Fig. S1A**. Scale bars = 1 cm.

**Fig. S2.**
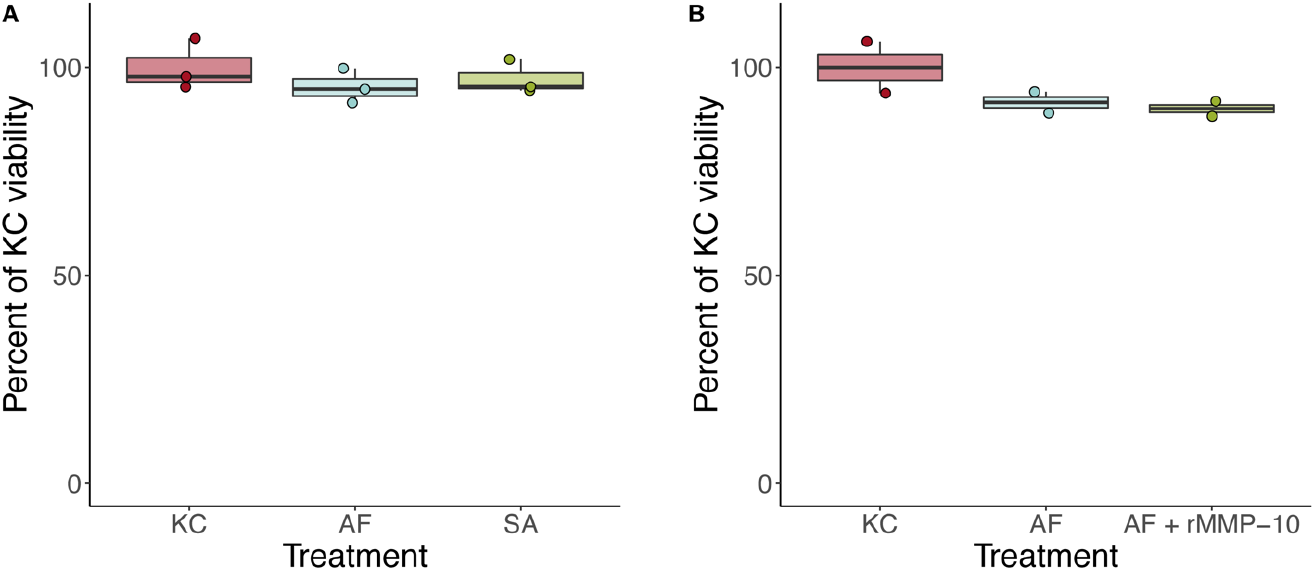
Viability of diabetic keratinocytes treated with bacterial conditioned media. (**A, B**) Viability of diabetic murine keratinocytes as measured by the alamarBlue assay. Data shown was normalized to the mean viability of the vehicle-treated wells (KC). N = 3 wells/condition (**A**), and 2 wells condition (**B**); data representative of 2 independent experiments.

**Fig. S3.**
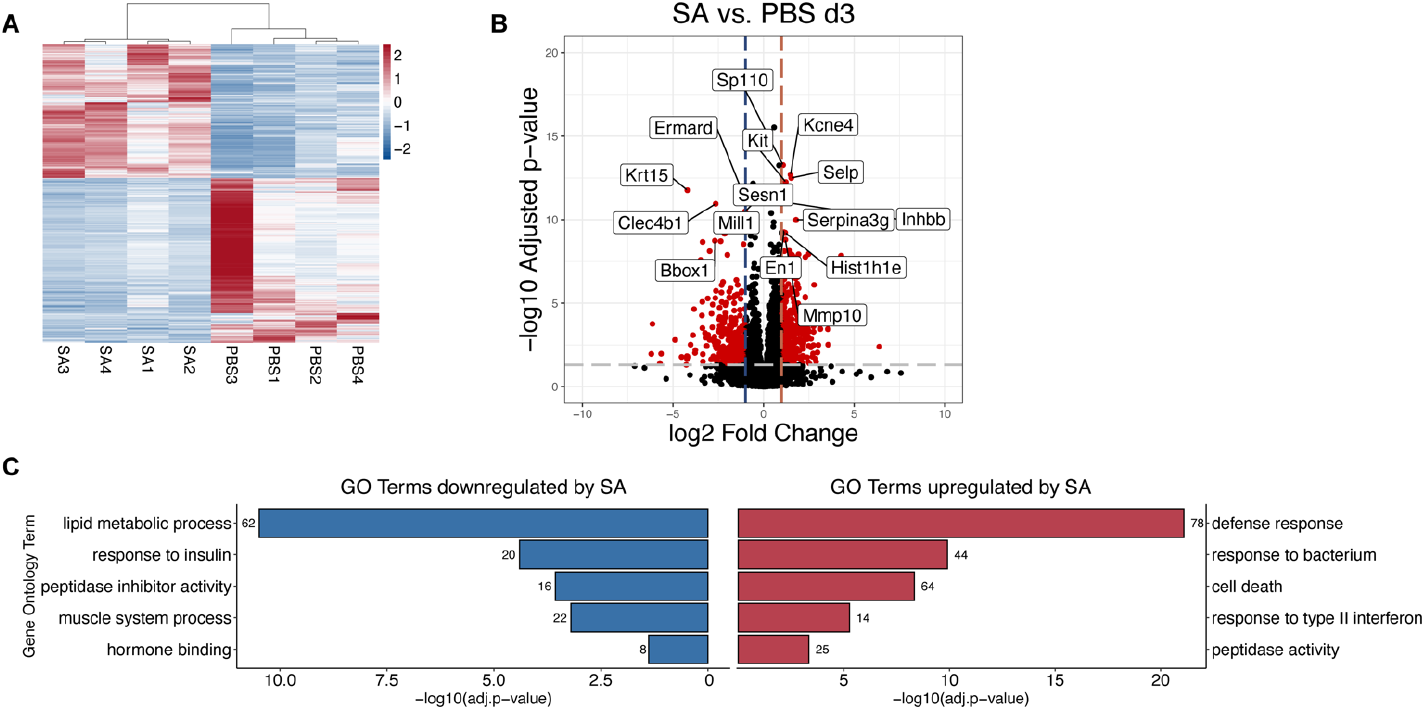
*S. aureus* wound treatment induces infection response transcriptional signatures. (**A**) Heatmap of differential expressed genes (DEG with FDR < 0.01) between the SA and PBS groups at day 3. Unsupervised clustering using Spearman’s correlation was used for hierarchical clustering of the samples, while Pearson’s correlation was used for gene clustering (heatmap rows, dendrogram not shown). Colors are used to codify log fold change values, with red values for increased expression and blue values for decreased expression. (**B**) Volcano plot demonstrating differentially expressed genes in SA vs. PBS wounds at day 3. Horizontal dashed line represents a cut off for the FDR p-adjusted value of <0.05. Vertical lines denote a fold change of ±2-fold. Genes that surpass both the p-value and fold change cut off are marked in red, with genes that are downregulated by SA to the left of the plot and upregulated to the right. The top 15 significant genes are highlighted with gene names on the plot. (**C**) Gene ontology (GO) enrichment was performed on differentially expressed genes for SA vs. PBS at d3. Up and downregulated GO terms of interest are highlighted, with the number of genes matching the term adjacent to the bar plot.

**Fig. S4.**
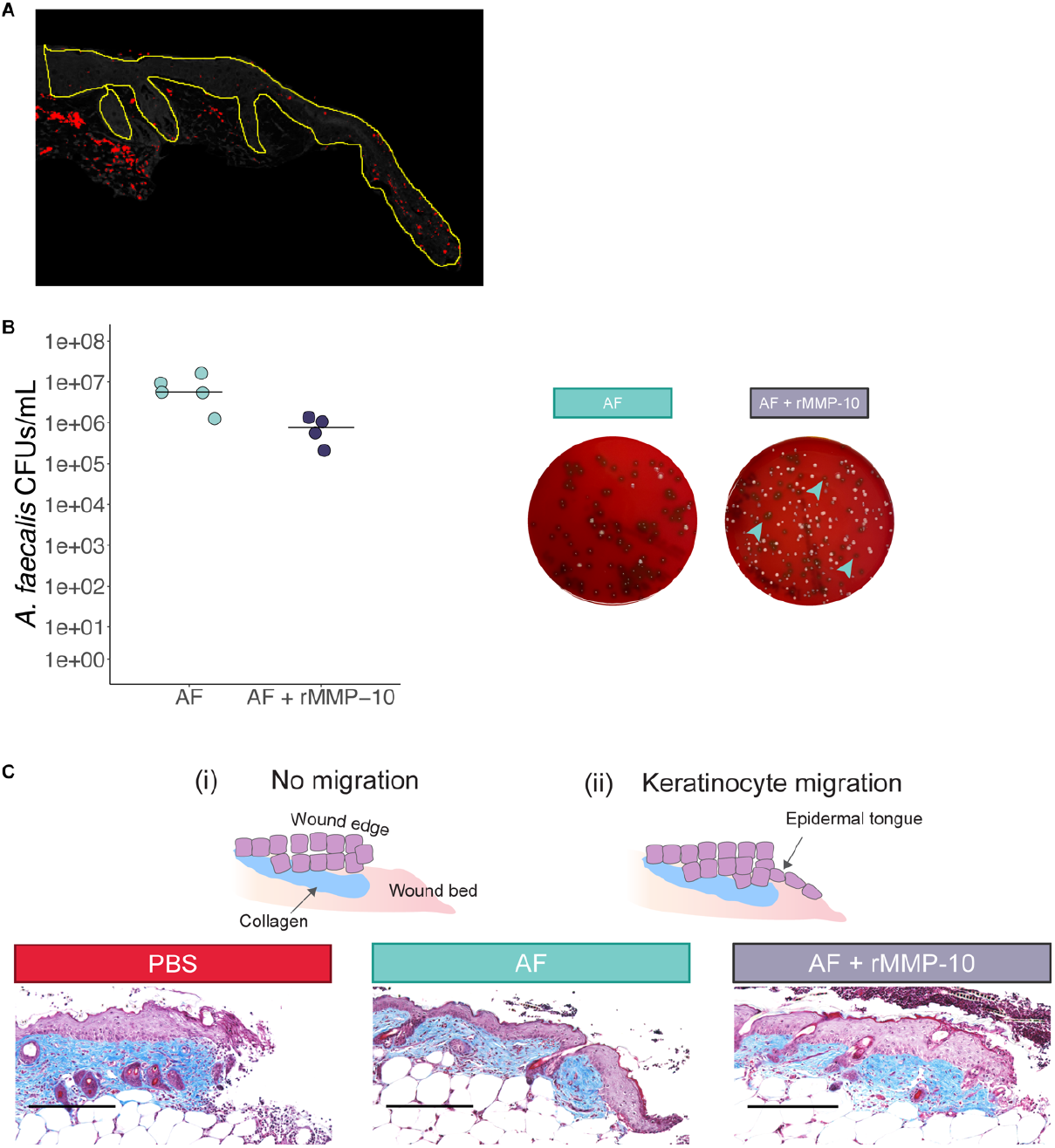
MMP-10 protein quantification and effects on *A. faecalis* quantification. (**A**) Sample representation of MMP-10 quantification using ImageJ. Shown is a wound image with thresholding performed so that positive pixels are red. The epidermal selection where quantification was performed is outlined in yellow. (**B**) Bacterial CFU quantification of day 3 wounds shown in **Fig. 4E**. On the right, representative blood agar plates from wound bacterial cultures are shown. The AF group is predominated by grey *A. faecalis* colonies. In the AF + rMMP-10 group, blue arrows point to examples of *A. faecalis* isolates; white colonies are the murine skin commensal *Staphylococcus xylosus* (colony identities confirmed by 16s rRNA gene sequencing). n of 5 mice per group for AF, 4 per group for AF + rMMP-10 and 1 wound per mouse. (**C**) Masson’s Trichrome staining of wound sections shown in **Fig. 4E**. The epidermis is stained in purple, and underlying collagen-rich dermis is blue, allowing for clear visualization of re-epithelialization at the wound tongue (i, ii). Representative images of each treatment group are shown. Scale bar = 250 μm.

